# A nutrient-derived dietary gradient links gut microbiome composition, network structure, and host physiology

**DOI:** 10.64898/2026.07.24.740531

**Authors:** Jiyan Xu, Maya Riabchenko, Theda U.P. Bartolomaeus, Alexander Müller, Luciana Hannibal, Roman Huber, Michael Jeitler, Víctor Hugo Jarquín-Díaz, Sofia K. Forslund-Startceva, Maximilian Andreas Storz

## Abstract

Diet is an important determinant of gut microbiome composition and host physiology. However, dietary exposure is often represented using categorical groups that may not fully capture variation in nutrient intake or its associations with gut microbiome and host biomarkers. Here, we analyzed diet-microbiome-host physiology relationships in healthy adults with long-term adherence (≥2 years) to omnivore, vegetarian, or vegan diets. We integrated four-day weighed dietary records, blood biomarkers, and 16S rRNA gene sequencing to characterize dietary intake, host physiology, and gut microbiome composition. We derived nutrient intake gradients by dimensionality reduction of representative nutrients and identified the second nutrient intake gradient (NIG2) as a quantitative axis reflecting variation between animal- and plant-leaning nutrient profiles, broadly recapitulating categorical dietary groups. While NIG2 showed no association with microbiome within-sample diversity, it explained gut microbiome compositional variation that was not detected using categorical dietary groups. Mediation analyses identified nominal candidate microbiome-mediated diet-biomarker relationships involving thyroid-related, hematological, and vitamin-related biomarkers. Network analyses of NIG2-defined animal-leaning and plant-leaning subsets identified subset-specific microbial associations and differences in microbial community structure and potential interactions. Overall, this continuous nutrient-derived dietary gradient provides a quantitative exposure metric for characterizing diet-microbiome-host physiology relationships.

**Graphical Abstract:** 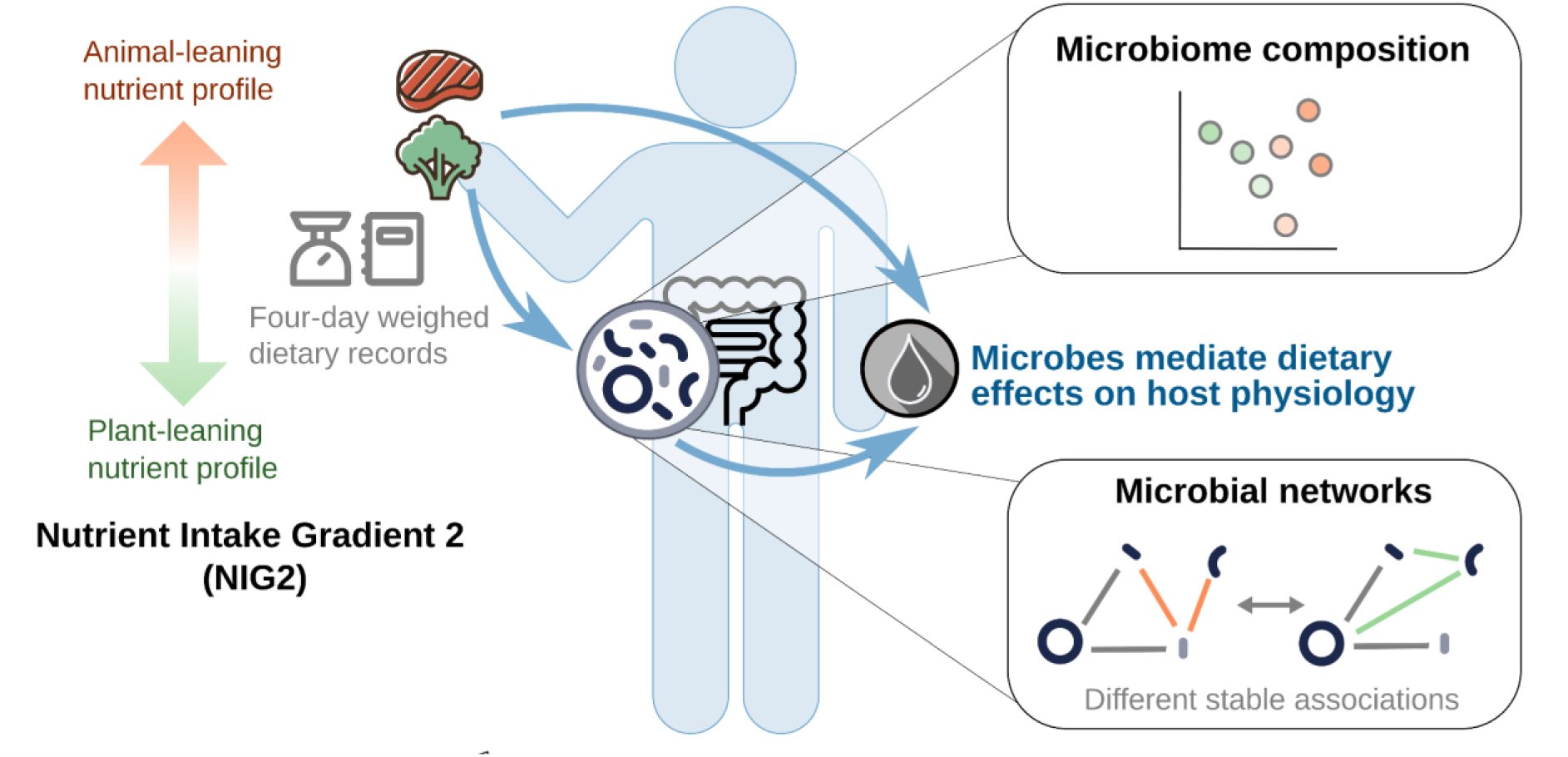

## Introduction

Diet and the gut microbiome are closely linked determinants of human physiology^1^. In particular, comparisons between dietary patterns have revealed consistent diet-associated variation in gut microbiome composition^2,3^. Thus, dietary intake patterns are relevant contributors to inter-individual variation in the gut microbiome. However, dietary group classifications represent broad behavioral categories and may not fully capture inter-individual variation in actual nutrient exposure, even among individuals adhering to the same dietary group^4,5^. Food choices, nutrient composition, and supplement use can vary substantially within dietary groups, potentially resulting in heterogeneous dietary exposures despite a shared dietary category. Consequently, direct nutrient-level measurements may offer a more comprehensive representation of dietary exposures relevant to detecting granular associations with the gut microbiome and host physiology, which may otherwise be obscured by general categorization^6^. At the same time, nutrients are not consumed in isolation but as correlated combinations reflecting broader dietary patterns^7^. Approaches that summarize correlated nutrient patterns may provide complementary information beyond analyses based on individual nutrients or categorical dietary groups.

The gut microbiome is associated with multiple aspects of host physiology, including metabolic^8^, hematological^9^, inflammatory^10^, and nutritional biomarkers^11^. Also, the microbiome has been proposed as a potential intermediate layer connecting diet and host phenotypes^12^. However, the extent to which microbiome variation reflects diet-associated host biomarker variation remains partially understood, particularly in healthy dietary cohorts.

The gut microbiome functions as a complex ecological community rather than a collection of independent taxa^13^. Microbial taxa are linked through shared metabolic pathways, resource competition, cross-feeding, and other ecological processes that are impacted or promoted by environmental processes^14^. To decipher subtle changes at the microbial community level promoted by nutrient-level dietary variation, it is necessary to characterize community organization and identify patterns of microbial relationships that are not apparent from abundance profiles alone^15^.

Despite efforts to identify diet-microbiome relationships, the use of categorical diet classifications may not provide enough resolution, especially in small cohorts to detect them. This becomes relevant when the aim is to understand the impact of diet-microbiome relationships in host health and physiology. Few studies have integrated nutrient-level dietary variation, microbiome structure and composition, and host biomarkers, within the same framework.

In this study, we investigated whether nutrient-derived dietary gradients could capture microbiome-relevant dietary variation across omnivore, vegetarian, and vegan individuals. We examined associations between nutrient-derived dietary gradients, gut microbiome composition, host blood biomarkers, and microbial association networks within a well-characterized healthy dietary cohort. Using an integrative approach, we aimed to evaluate how nutrient-associated dietary variation is reflected across multiple levels of microbiome organization and host physiology.

## Methods

### Recruitment, cohort characterization, and stool sample collection

Healthy adults adhering to a long-term omnivore, vegetarian or vegan dietary pattern (≥ 2 years) were recruited in Freiburg, Germany. Dietary pattern assignment followed established definitions^16^ and was based on self-identification in combination with weighed food records, providing high-resolution estimates of average daily nutrient intakes (ADI). The formula to estimate ADIs has been discussed elsewhere in detail^17,18^. Demographic metadata, supplement intake and blood samples were collected as previously described^19–22^. In this cross-sectional observational study, one stool sample was collected from each participant after recruitment. Stool samples were preserved and aliquoted in separate 1.5-mL cryovials and stored at −80 °C until measurement.

For cohort characterization, differences in demographic and clinical characteristics across dietary groups were assessed using chi-square tests for categorical variables and Kruskal–Wallis tests for continuous variables. Nominal *p* values are reported for these descriptive comparisons.

### DNA extraction, amplification and sequencing

DNA extraction was performed using the ZymoBIOMICS 96 MagBead DNA Kit (Zymo Research, Irvine, CA, USA) according to the manufacturer’s protocol. Amplification of the 16S rRNA gene targeted the V3-V4 hypervariable regions using the Quick-16S Primer Set V3-V4 (Zymo Research Europe, Freiburg, Germany). Sequencing libraries were prepared and sequenced on the Illumina MiSeq platform using the MiSeq Reagent Kit V3 (600 cycles), generating paired-end reads.

### 16S rRNA gene sequence preprocessing and taxonomic profiling

Sequencing reads were processed using LotuS2 (v.2.34)^23^ with the SILVA 138.1 database for taxonomic classification. Host-derived reads were filtered using the human reference genome hg38.p14 to remove off-target sequences. Operational taxonomic units (OTUs) unassigned at the phylum level were excluded. Prior to rarefaction, two samples with sequencing depth below 36,887 reads were excluded. The remaining OTU table was rarefied to an equal sequencing depth of 36,887 reads per sample using the rtk package (v.0.2.6.1)^24^. The rarefied OTU table was integrated with the sample metadata and OTU taxonomy into a single object using phyloseq (v.1.48.0)^25^. For further analyses, counts were aggregated to the genus level using the *tax_glom* function from phyloseq.

### Nutrient intake feature summarization

To characterize dietary patterns beyond categorical dietary group labels, we analyzed nutrient intake from food across omnivore, vegetarian, and vegan individuals. For each nutrient, a linear regression model was fitted with nutrient intake as the response variable and dietary group as the main explanatory variable, adjusting for host covariates (sex, age, and body mass index (BMI)), using the stats package (R v.4.4.3)^26^. The diet-attributable proportion of explained variance was quantified as the partial R² comparing the full model, including dietary group and covariates, with a reduced model including covariates only. The *p* values for dietary group effects were adjusted for multiple testing using the Benjamini–Hochberg procedure^27^. Nutrients with FDR-adjusted *p* < 0.1 were considered diet-associated for nutrient screening.

To account for redundancy among nutrients due to co-consumption and shared dietary sources, nutrients were hierarchically clustered based on pairwise Spearman correlations of intake values across individuals. Clusters were defined using a correlation threshold of |ρ| ≥ 0.7. Within each cluster, the nutrient with the largest diet-attributable partial R² was selected as the representative food-derived nutrient for downstream analysis.

Principal component analysis (PCA) was performed on the selected representative food-derived nutrients to summarize multivariate nutrient intake variation across individuals. Nutrient intake values were centered and z-score standardized prior to PCA. The first two principal components were retained based on the scree plot and are hereafter referred to as nutrient intake gradient 1 (NIG1) and nutrient intake gradient 2 (NIG2). Unless otherwise stated, NIG1 and NIG2 refer to the primary gradients obtained exclusively from food-derived nutrient intake. Individual PCA scores and nutrient loading vectors for the first two components were extracted for visualization and interpretation. Nutrient contributions to each gradient were calculated from the PCA loadings, and the quality of representation of each nutrient on the NIG1-NIG2 plane was quantified as the cumulative cos² across the first two principal components. For visualization only, nutrient loading vectors were multiplied by a fixed scaling factor of 4; this transformation did not affect the PCA results or the interpretation of the underlying loadings. Differences in individual PCA scores between dietary groups were assessed using one-way analysis of variance (ANOVA), followed by Tukey’s honestly significant difference (HSD) post hoc test to identify significant pairwise differences. Model assumptions were evaluated before inference. Normality of residuals was assessed using the Shapiro–Wilk test together with visual inspection of Q–Q plots, and homogeneity of variances was assessed using Levene’s test with the car package (v.3.1-3)^28^. Supporting analyses were performed to assess associations between nutrient intake gradients and demographic covariates. Sex differences were tested using Wilcoxon rank-sum tests, while associations with age and BMI were tested using Spearman correlation. Linear regression lines with 95% confidence intervals were shown for visualization.

As a sensitivity analysis to evaluate potential effects of non-food-derived nutrients, the same PCA workflow was repeated using total nutrient intake, including food-derived intake and available supplement-derived intake. The resulting axes are referred to as total-intake NIG1 and total-intake NIG2. Nutrient variables, scaling parameters, and PCA loadings used to define the food-derived and total-intake nutrient intake gradients are provided in Supplementary Table 1. Concordance between the food-derived and total-intake PCA configurations was assessed using Procrustes analysis based on the first two PCA axes. Agreement between food-derived NIG2 and total-intake NIG2 scores was further evaluated using Pearson correlation.

### Microbiome diversity and composition analyses

Alpha diversity was quantified at the genus level using Shannon, Simpson, and Chao1 indices calculated with the *estimate_richness* function in the phyloseq package. Associations between alpha diversity and NIG2 were assessed using linear regression models implemented with the stats package (v.4.4.3), adjusting for host covariates. Model assumptions, including linearity, normality of residuals, and homoscedasticity, were evaluated using diagnostic plots with the car package (v.3.1-3)^28^. Microbial community composition was assessed using Bray-Curtis dissimilarity calculated from genus-level relative abundance profiles, with the vegdist function in the vegan package (v.2.7-2)^29^. Principal coordinate analysis (PCoA) was used to visualise differences in overall microbiome composition along the NIG2. Statistical associations between microbial community composition and NIG2 were tested using permutational multivariate analysis of variance (PERMANOVA) implemented in the *adonis2* function of the vegan package, with marginal effects specified (by = “margin”) and 9,999 permutations. Models included NIG2 as the main explanatory variable and also included sex, age, and BMI as covariates. To quantify the association between NIG2 and the main axes of microbiome variation, linear regression models were fitted with PCoA1 or PCoA2 as the response variable and NIG2 as the main predictor, adjusting for sex, age, and BMI. Partial R² values were calculated to quantify the variance attributable to each predictor after accounting for covariates. To identify genera associated with the first two ordination axes, genus-level abundance vectors were fitted onto the first two PCoA axes using the *envfit* function in vegan with 999 permutations. The *p* values were adjusted using the Benjamini–Hochberg procedure, and genera with FDR-adjusted *p* < 0.1 were considered associated with the ordination space. For visualization, the ten genera with the largest envfit R² among genera with FDR-adjusted *p* < 0.1 were displayed as vectors. Vectors were scaled by a fixed factor 0.5 for visualization; this scaling did not affect the underlying envfit statistics.

As a sensitivity analysis, PERMANOVA was repeated using total-intake NIG2 instead of food-derived NIG2 to assess whether the association between nutrient intake gradients and microbiome composition was robust to the inclusion of supplement-derived nutrient intake.

### Regression-based mediation analyses of microbial genera

To investigate whether gut microbial features may mediate associations between dietary nutrient patterns and host blood biomarkers, we performed regression-based mediation analyses using NIG2 as the exposure, microbial genera as candidate mediators, and blood biomarkers as outcomes. Microbial genera were filtered before analysis to reduce sparsity and improve statistical robustness. Genera with prevalence ≥10% and mean relative abundance ≥1 × 10⁻⁴ across samples were retained. To reduce redundancy among blood biomarkers, biomarkers were grouped into physiologically defined modules based on clinical and biological relevance and subsequently clustered within modules using pairwise Spearman correlations across individuals. Clusters were defined using a correlation threshold of |ρ| ≥ 0.7. Within each cluster, one representative biomarker was selected based on biological relevance and interpretability, prioritizing the most direct physiological measurement over derived variables. The hierarchical clustering and selected representative biomarkers are shown in Supplementary Fig. S1, while detailed biomarker information and representative selections are provided in Supplementary Table S1. Microbial genus relative abundances and blood biomarker values were log-transformed using *log1p* to reduce skewness and stabilize variance. The distribution of NIG2 was assessed by visual inspection and by calculating the sample skewness statistic using the moments package (v.0.14.1)^30^. The observed skewness (−0.32) indicated no pronounced skewness (Supplementary Fig. S2). Therefore, NIG2 was not log-transformed prior to standardization. NIG2, microbial abundances, and blood biomarkers were then standardized to zero mean and unit variance, allowing regression coefficients to be compared across models.

The mediation workflow first evaluated three regression paths: path a tested associations between NIG2 and microbial abundance, path b tested associations between microbial abundance and blood biomarkers, and path c tested associations between NIG2 and blood biomarkers. All three regression models were adjusted for host covariates, using the stats package. For each microbial genus or biomarker, regression coefficients and *p* values were estimated. To reduce multiple testing and focus subsequent mediation testing on biologically plausible candidate relationships, microbial genera nominally associated with NIG2 in path a and biomarkers nominally associated with NIG2 in path c were retained as candidate mediators and outcomes, respectively. Path b associations were then evaluated for all retained genus-biomarker combinations.

Formal mediation analyses were performed in the forward direction (NIG2 to genera to biomarkers) using the mediation package (v.4.5.0)^31^. For each candidate genus-biomarker pair, the average causal mediation effect (ACME), average direct effect (ADE), total effect, and proportion mediated were estimated. Confidence intervals and *p* values were obtained using nonparametric bootstrap resampling with 1,000 simulations. A mediation pathway was considered nominally significant when both the ACME and the proportion mediated were significant at *p* < 0.05.

Reverse-direction mediation analyses were additionally performed to assess the directionality of the observed genus-biomarker relationships. In these models, blood biomarkers were modeled as candidate mediators between NIG2 and microbial genera, using the same regression-based framework, covariate adjustment, bootstrap procedure, and significance criteria as in the forward mediation analyses.

### Conditional microbial association network inference and stability-based edge comparison

To investigate whether microbial association structure differed along the dietary nutrient gradient, participants were stratified by the median value of NIG2. Individuals with NIG2 values above the median were classified as animal-leaning, and individuals with NIG2 values below the median were classified as plant-leaning. This median-based stratification generated two subsets of comparable sample size (n= 52) for downstream network inference.

Microbial networks were inferred using a metadata-adjusted Gaussian graphical model framework. A filtering criteria consistent with the one used in the mediation analyses was implemented. Genus-level abundance profiles were first transformed using a modified centered log-ratio transformation (mCLR) to account for compositionality and zero inflation using SPRING package (v.1.0.4)^32^. Host covariates were incorporated as additional nodes before network inference, with sex encoded as a binary variable, and both age and BMI as continuous covariates scaled to zero mean and unit variance. This framework allowed the estimation of conditional microbial associations while accounting for variation related to host covariates. A rank-based nonparanormal skeptic estimator was then applied using the *huge.npn* function with npn.func = “skeptic” from the huge package (v.1.3.5)^33^ to estimate a latent Gaussian copula correlation matrix. Graphical lasso was applied to the resulting correlation matrix using the glasso package (v.1.11)^34^ to estimate sparse conditional association networks.

The sparsity parameter λ was selected using stability-based model selection with the StARS (Stability Approach to Regularization Selection) criterion ^35,36^ implemented in the pulsar (v.0.3.11)^37^. The λ sequence was generated using lambda.min.ratio = 0.05 and nlambda = 30. Stability selection was performed using 50 repeated subsamples with a subsampling ratio of 0.8, a StARS instability threshold of 0.05, and a fixed random seed. The selected λ was then used for full-data network inference and repeated subsampling-based network estimation.

For downstream topological analyses and visualization, host covariates nodes were removed and analyses were restricted to the microbe-only subnetworks. Only positive microbial associations were retained. The largest connected component (LCC) of each inferred network was extracted for topological characterization. Network topology was summarized using density, LCC size, mean degree, and modularity, calculated with the igraph (v.2.1.4)^38,39^.

To compare edge stability between animal-leaning and plant-leaning subnetworks, networks were repeatedly inferred across 50 subsamples within each subset. Edge stability was defined as the proportion of subsampling networks in which a given edge was detected. Edges present in ≥90% of animal-leaning subsampling networks and ≤10% of plant-leaning subsampling networks were classified as animal-leaning stable edges. Plant-leaning stable edges were defined conversely. Edges present in ≥90% of subsampling networks in both subsets were classified as shared stable edges. Subset-differential stable edges were defined as edges classified as either animal-leaning or plant-leaning stable. Taxa were ranked by the number of subset-differential stable edges in which they participated, and the 15 most frequently involved taxa were selected for centrality and descriptive analyses. Eigenvector centrality was summarized across repeated subsampling networks and compared between animal-leaning, plant-leaning, and overall networks. Prevalence was defined as the proportion of samples in which a taxon was detected in the rarefied genus-level count table. Differences in prevalence between animal- and plant-leaning subsets were assessed using Fisher’s exact tests, followed by Benjamini–Hochberg correction. Relative abundance was calculated from rarefied genus-level counts and visualized on a log10-scaled percentage axis. Differences in relative abundance between animal- and plant-leaning subsets were tested using linear models fitted to log10-transformed relative abundance with adjustment for sex, age, and BMI, followed by Benjamini–Hochberg correction.

## Results

### General sequence processing statistics

The final decontaminated dataset used for the analysis contained 5,503,124 sequences (min. 36,887, max. 76,585) from 104 samples, with an average of 52,915 reads per sample, which passed the pre-processing and quality checks and were assigned to OTUs. After rarefaction, 724 OTUs were agglomerated into 191 genera and were included in all analyses.

### Cohort overview and demographic characteristics

After 16S rRNA sequencing quality control, 104 participants were retained for microbiome analysis, including 35 omnivores, 35 vegetarians, and 34 vegans (Fig. 1a). Sex distribution did not differ significantly across dietary groups (chi-square test, χ² = 1.21, df = 2, *p* = 0.55), and no statistically significant differences were detected for age (Kruskal–Wallis test, χ² = 0.71, df = 2, *p* = 0.70) or BMI (Kruskal–Wallis test, χ² = 4.56, df = 2, *p* = 0.102), indicating that the three dietary groups were broadly comparable with respect to these demographic characteristics.

**Fig. 1:**
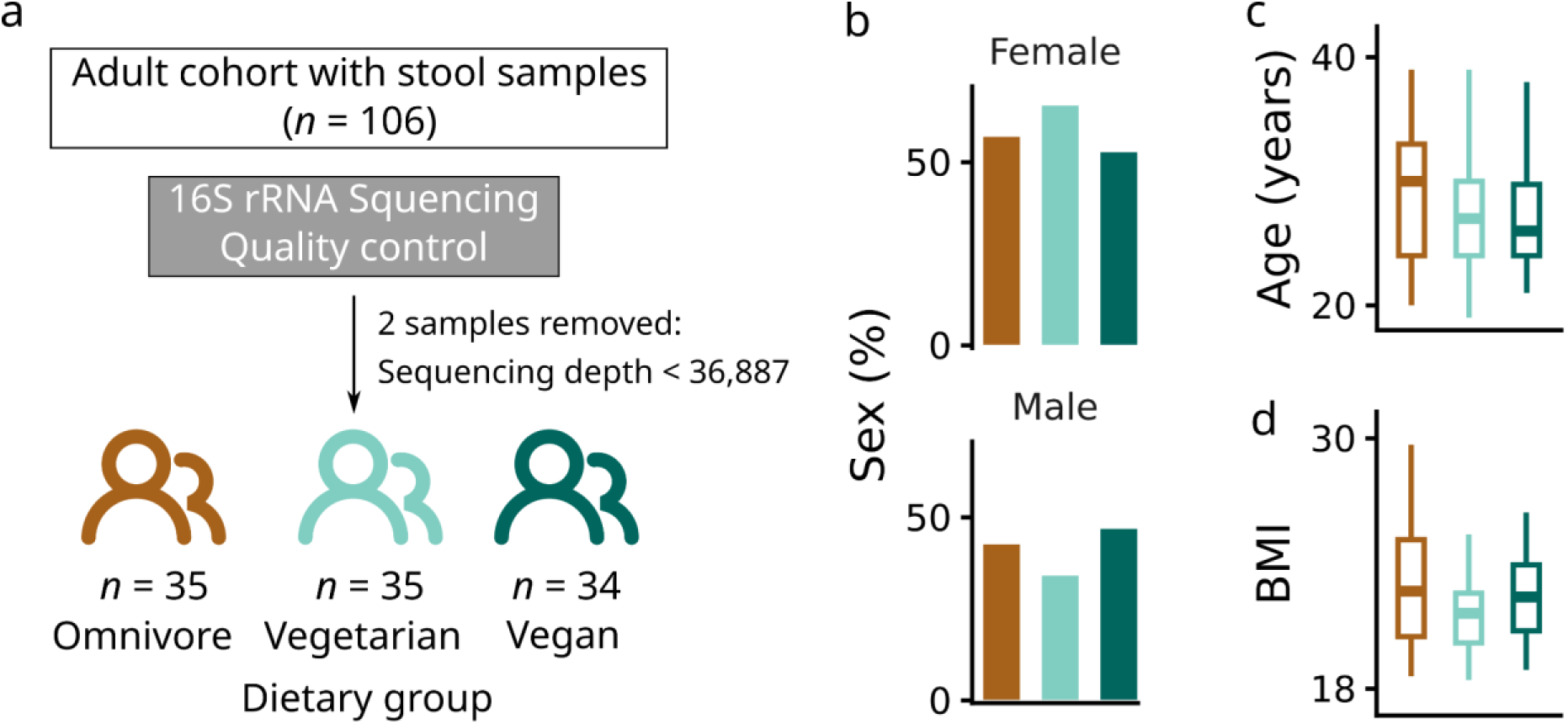
Study cohort and demographic characteristics. **a,** Overview of the study cohort and 16S rRNA sequencing quality control. Among 106 adults with available stool samples, two samples were removed after sequencing quality control, resulting in a final microbiome cohort of 104 participants, including 35 omnivores, 35 vegetarians, and 34 vegans. No significant group differences were observed for sex distribution (**b**), age (**c**), or BMI (**d**) across dietary groups; sex was assessed using a chi-square test, and age and BMI were assessed using Kruskal–Wallis tests.

### A nutrient intake gradient captures dietary variation across groups

Dietary group explained substantial variation in the intake of multiple nutrients after adjustment for host covariates (Fig. 2a). Nutrients commonly linked to animal-derived foods and lipid intake, including vitamin B12, vitamin D, and saturated fatty acids (SFA), ranked among the nutrients with the highest diet-attributable explained variance (partial R² > 0.20; Fig. 2a). Several nutrients typically associated with plant-based dietary intake, including fiber, folic acid, and vitamin C, showed smaller diet-attributable explained variance in nutrient-wise models, suggesting that plant-associated dietary variation was distributed across multiple correlated nutrients rather than dominated by a single nutrient feature. Hierarchical clustering of nutrient intake correlations (|ρ| ≥ 0.7) revealed groups of co-consumed nutrients, consistent with structured dietary patterns rather than isolated nutrient effects. To reduce redundancy while retaining diet-relevant information, one representative nutrient was selected from each correlation-defined cluster based on the magnitude of diet-attributable explained variance (Fig. 2a).

**Fig. 2:**
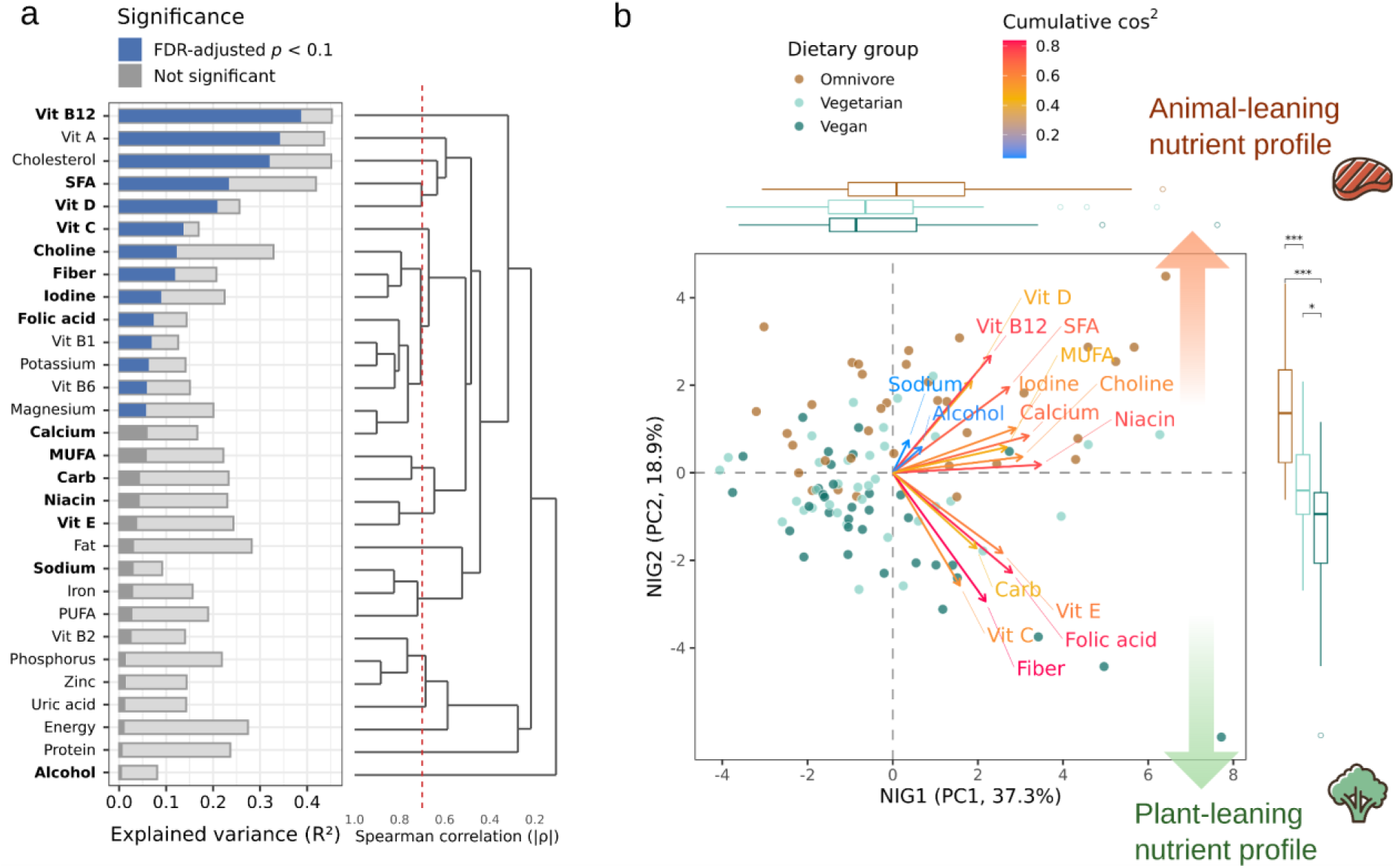
A nutrient intake gradient captures diet-associated variation in nutrient profiles. **a,** Diet-associated variation in nutrient intake. For each nutrient, the proportion of variance explained by the dietary group was estimated using linear models adjusted for sex, age, and BMI. Light grey bars indicate total model R², while blue or dark grey segments represent the partial R² attributable to dietary groups. Nutrients with significant diet effects after Benjamini–Hochberg correction are shown in blue (FDR-adjusted *p* < 0.1); non-significant diet effects are shown in grey. The dendrogram represents the hierarchical clustering of the nutrients based on Spearman correlations of intake values across individuals. Nutrients were grouped using a correlation threshold of |ρ| ≥ 0.7, indicated by the red dashed line. Within each correlation-defined cluster, the nutrient with the largest diet-attributable partial R² was selected as a representative nutrient for downstream dimensionality reduction and is shown in bold. **b,** Principal component analysis (PCA) of the selected representative nutrients. Each point represents an individual nutrient profile and is colored by dietary group. Nutrient arrows indicate loading vectors on the first two principal components, hereafter referred to as nutrient intake gradient 1 (NIG1) and nutrient intake gradient 2 (NIG2). Arrow direction reflects the association of each nutrient with the ordination axes, whereas arrow color denotes the quality of representation on the NIG1-NIG2 plane, quantified as the cumulative cos² across both axes. For visualization, arrow lengths were multiplied by a fixed factor of 4. Marginal boxplots show the distribution of individual scores along NIG1 and NIG2 by dietary group. Positive NIG2 values correspond to an animal-leaning nutrient profile, whereas negative NIG2 values correspond to a plant-leaning nutrient profile. Statistical significance in marginal boxplots was assessed using one-way ANOVA followed by Tukey’s honestly significant difference post hoc test. *Adjusted *p* < 0.05; ***adjusted *p* < 0.001.

Principal component analysis of these representative nutrients identified two major axes, defined as nutrient intake gradient 1 (NIG1) and nutrient intake gradient 2 (NIG2), explaining 37.3% and 18.9% of the variance, respectively (Fig. 2b, Supplementary Fig. S3a). NIG1 captured broad variation across multiple nutrients but did not clearly separate dietary groups. In contrast, NIG2 captured a diet-associated nutrient gradient, with positive loadings for animal- and lipid-associated nutrients, including vitamin B12, vitamin D, and SFA, and negative loadings for plant-associated nutrients, including fiber, vitamin C, and folic acid (Fig. 2b, Supplementary Fig. S3b-e). Consistent with this loading structure, individual NIG2 scores recapitulated the categorical dietary groups (omnivore, vegetarian, and vegan) from the participants along a continuous dietary gradient (Fig. 2b). NIG2 scores were highest in omnivores and progressively lower in vegetarian and vegan participants (ANOVA, F(2,101)= 34.26, *p*= 4.4×10⁻¹²), with significant pairwise differences between all dietary groups (Tukey HSD, all adjusted *p* < 0.01). Nutrient loading vectors in the PCA biplot showed opposing contributions from animal- and plant-associated nutrients along NIG2, supporting its interpretation as an animal- to plant-leaning nutrient intake gradient. NIG2 was therefore used as the primary continuous dietary exposure in downstream microbiome and mediation analyses.

Associations between both NIGs and host covariates were further examined. NIG1 was associated with all host covariates, whereas NIG2 was only associated with age but not with sex or BMI (Supplementary Fig. S4). Subsequent regression analyses therefore included relevant host covariates to account for potential confounding.

Because some participants reported supplement use, we repeated the nutrient PCA using total nutrient intake, including supplement-derived intake where available. The total-intake PCA preserved the diet-associated separation along the second axis, with omnivore, vegetarian, and vegan participants remaining significantly separated along total-intake NIG2 (ANOVA, F(2,101)= 27.9, *p*= 2.26 × 10^−10^; Supplementary Fig. S5a). Food-derived and total-intake PCA configurations were highly concordant by Procrustes analysis (r = 0.91, *p* = 1.00 × 10^−4^), and food-derived NIG2 scores were strongly correlated with total-intake NIG2 scores (Pearson r = 0.84, *p* = 2.95 × 10^−29^; Supplementary Fig. S5b,c). These sensitivity analyses supported the robustness of the food-derived NIG2 used as the primary dietary exposure in downstream analyses.

### NIG2 explains gut microbiome compositional variation

NIG2 has no significant association with gut microbiome alpha diversity metrics after adjustment for host covariates (Supplementary Fig. S6). However, gut microbial community composition varied continuously along NIG2. In the Bray–Curtis PCoA ordination, samples with higher NIG2 values tended to be positioned toward positive values of the first principal coordinate, whereas samples with lower NIG2 values were positioned toward negative values (Fig. 3a). PERMANOVA confirmed a significant association between NIG2 and overall microbial composition after adjustment for host covariates (R² = 0.036, *p* = 0.002). BMI was also significantly associated with microbial composition, although with a smaller effect size (R² = 0.023, *p* = 0.016). In contrast, the categorical classification of diet did not detect the changes in microbial composition (Supplementary Fig. S7; PERMANOVA, R² = 0.024, *p* = 0.212). Consistent with the ordination pattern, linear regression analysis showed that NIG2 was positively associated with PCoA1 after adjustment for host covariates (β = 0.028, partial R² = 0.111, *p* = 6.58 × 10^−4^; Fig. 3b). BMI also showed a smaller positive association with PCoA1 (β = 0.010, partial R² = 0.038, *p* = 0.049). No significant association was observed for the second principal coordinate (PCoA2). These results indicate that the primary axis of genus-level microbiome compositional variation was aligned with the animal- to plant-leaning nutrient intake gradient. Genus vectors fitted onto the PCoA ordination using envfit further illustrated the taxonomic structure underlying this compositional variation (Fig. 3a). *Prevotella* 9 was oriented toward the plant-leaning end of the NIG2-associated PCoA1 axis, whereas *Bacteroides* and *Blautia* were oriented toward the animal-leaning end. Other fitted taxa, including the Christensenellaceae R-7 group and unclassified Ruminococcaceae, were oriented mainly along PCoA2 rather than along the NIG2-associated PCoA1 axis.

**Fig. 3:**
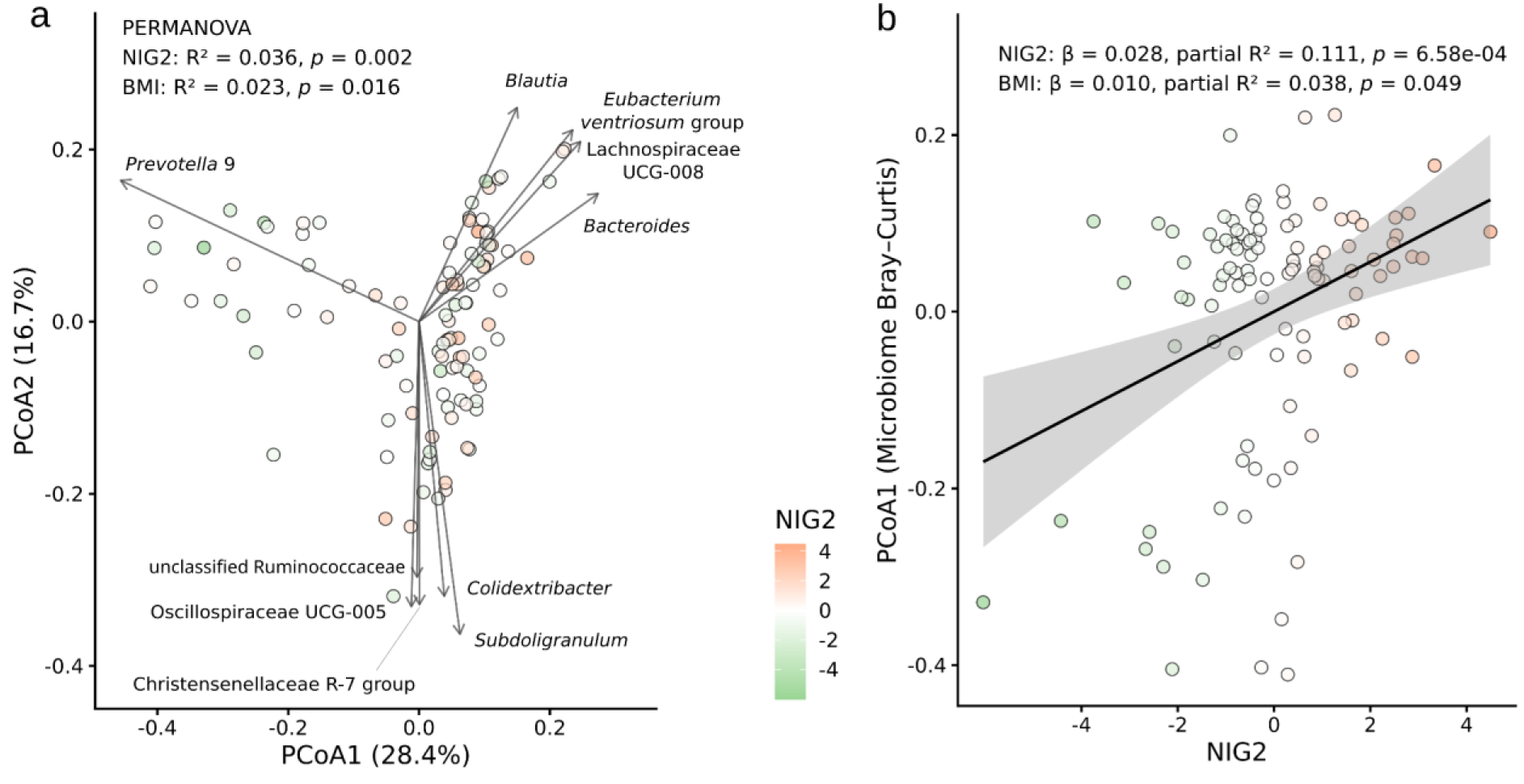
NIG2 explains gut microbiome compositional variation. **a,** Principal coordinate analysis (PCoA) of Bray-Curtis dissimilarities based on genus-level microbial composition. Each point represents one sample and is colored by NIG2, the continuous dietary gradient derived from food-derived nutrient intake profiles. Selected genus vectors indicate the ten FDR-significant genera with the strongest associations with the PCoA ordination based on envfit; vector direction indicates the direction of increasing genus abundance in the ordination space. The association between NIG2 and overall microbial composition was tested using PERMANOVA adjusted for sex, age, and BMI (R² = 0.036, *p* = 0.002). BMI was also associated with microbial composition with a smaller effect size (R² = 0.023, *p* = 0.017). **b,** Association between NIG2 and the first principal coordinate of microbial composition. Points represent individual samples and are colored by NIG2. The solid line indicates the fitted linear regression, and the shaded area indicates the 95% confidence interval. NIG2 was positively associated with PCoA1 after adjustment for sex, age, and BMI (β = 0.028, partial R² = 0.111, *p* = 6.58 × 10^−4^). BMI also showed a smaller positive association with PCoA1 (β = 0.010, partial R² = 0.038, *p* = 0.049).

As an additional sensitivity analysis, total-intake NIG2 remained significantly associated with genus-level microbiome composition, although with a smaller effect size than food-derived NIG2 (PERMANOVA, R² = 0.019, *p* = 0.048) indicating the low effect introduced by supplements.

### Gut microbial genera mediate dietary effects on host physiology

We applied a regression-based mediation framework to investigate candidate diet-microbiome-host physiology relationships. We used NIG2 as the measurement of diet exposure, microbial genera as candidate mediators, and blood biomarkers as outcomes representing impact in host physiology (Fig. 4a). The mediation framework comprised three association paths: path a between NIG2 and microbial abundance, path b between microbial abundance and blood biomarkers, and path c between NIG2 and blood biomarkers. After filtering, the mediation analysis included 108 microbial genera and 43 blood biomarkers. In the regression-based screening analysis, NIG2 was nominally associated with the abundance of 15 microbial genera in path a and with 8 blood biomarkers in path c, representing the total association between NIG2 and host physiology (Fig. 4b). Path c associations involved lipid-, hematological-, thyroid-, and vitamin-related biomarkers, indicating that nutrient-level dietary variation was associated with multiple physiological systems.

**Fig. 4:**
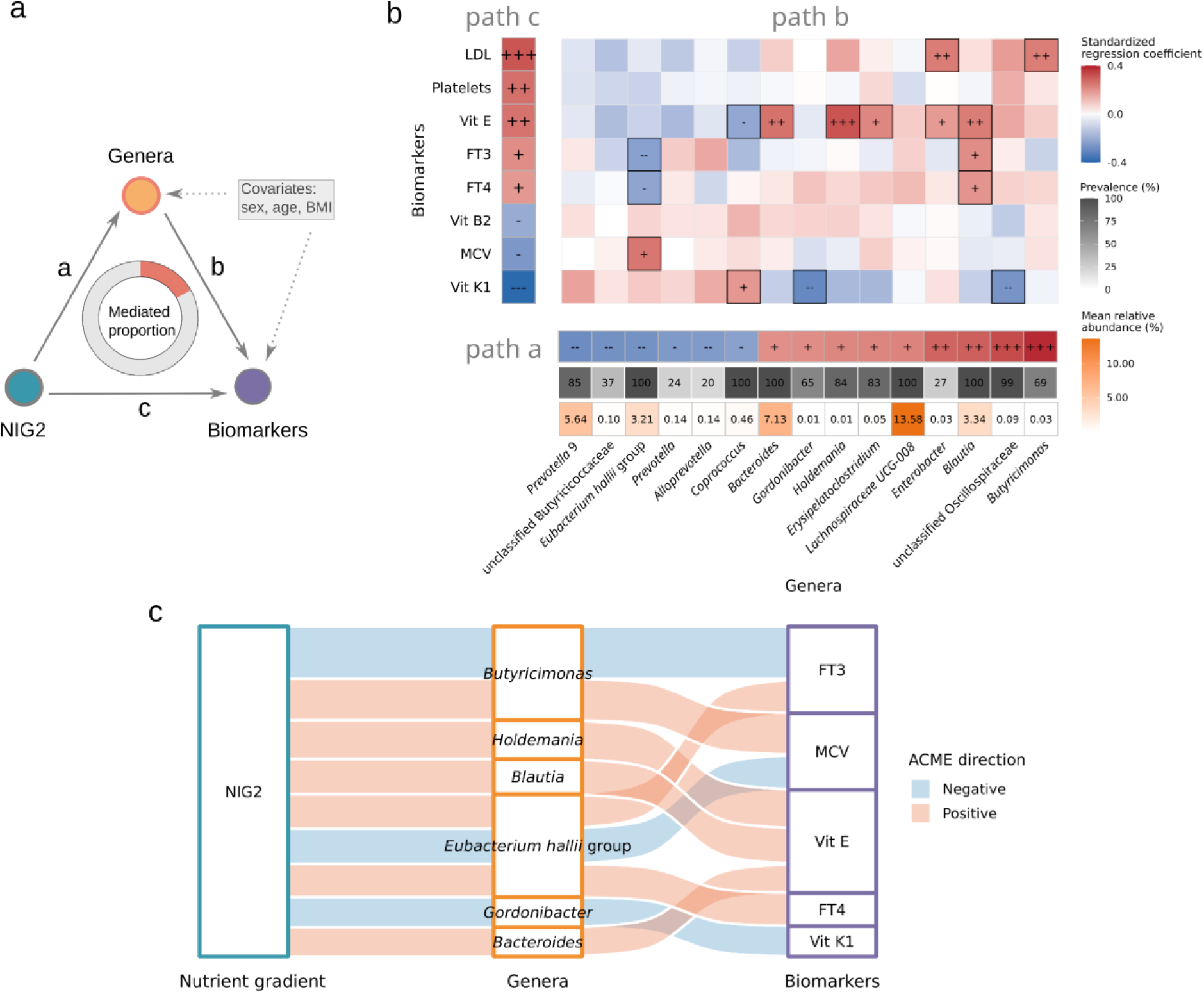
Regression-based mediation analysis linking NIG2, gut microbial genera, and blood biomarkers. **a,** Conceptual framework of the mediation analysis. NIG2 was modeled as the exposure variable, microbial genera as candidate mediators, and blood biomarkers as outcomes. Path a represents the association between NIG2 and microbial genera, path b represents the association between microbial genera and blood biomarkers, and path c represents the total association between NIG2 and blood biomarkers. All regression models were adjusted for sex, age, and BMI. **b,** Regression-based screening of associations used to define candidate mediation relationships. The bottom horizontal heatmap shows path a associations between NIG2 and microbial genera. The left vertical heatmap shows path c associations between NIG2 and blood biomarkers. The main heatmap shows path b associations between microbial genera and blood biomarkers for candidate genera and biomarkers supported by nominally significant path a and path c associations. Tile color represents standardized regression coefficients. Symbols indicate nominal significance levels, with symbol direction reflecting the sign of the coefficient: “+++” or “---”, *p* < 0.001; “++” or “--”, *p* < 0.01; “+” or “−”, *p* < 0.05. The two annotation rows below the path a heatmap indicate genus prevalence and mean relative abundance across the study cohort. **c,** Nominally significant mediation relationships identified in the forward mediation analysis from NIG2 to microbial genera to blood biomarkers. Flow color indicates the direction of the average causal mediation effect (ACME), with blue indicating negative mediation effects and orange indicating positive mediation effects. Flow width is proportional to the absolute ACME estimate. Only mediation relationships with nominally significant ACME and proportion mediated are shown.

Using microbial genera and blood biomarkers supported by nominally significant path a and path c associations, forward mediation analyses identified nine candidate mediation relationships with significant average causal mediation effects (ACME) and proportions mediated (Fig. 4c; Supplementary Table S3). These candidate relationships involved *Bacteroides*, *Blautia*, *Eubacterium hallii* group, *Gordonibacter*, *Holdemania*, and *Butyricimonas*, linking NIG2-associated genera to thyroid-related biomarkers (FT3 and FT4), the hematological marker MCV, and the fat-soluble vitamins vitamin E and vitamin K1. Together, these candidate mediation relationships suggest that NIG2-associated changes in the gut microbiome may partially mediate associations between nutrient-level dietary variation and multiple aspects of host physiology. The involvement of multiple microbial genera and biomarker categories further supports the biological consistency of these candidate diet-microbiome-host relationships.

Reverse-direction mediation analyses, in which blood biomarkers were modeled as candidate mediators between NIG2 and microbial genera, recovered the same genus-biomarker combinations identified in the forward mediation analyses under the predefined nominal significance criteria (Supplementary Table S3). This consistency across mediation directions indicates that the observed relationships were reproducible within the applied mediation framework, although the directionality of these associations could not be resolved in this cross-sectional study. These candidate mediation relationships should be interpreted as hypothesis-generating as any remain significant after correction for multiple testing.

### Stability of microbial communities differed between NIG2-defined animal- and plant-leaning subsets

To assess whether microbial associations within the community structure differed along the animal-plant nutrient intake gradient, participants were stratified into animal-leaning and plant-leaning subsets according to the median NIG2 value (Fig. 5a). Sparse microbial conditional association networks were inferred separately within each subset using a metadata-adjusted Gaussian graphical model framework, with the sparsity parameter λ selected by StARS-based stability selection (Fig. 5b).

**Fig. 5:**
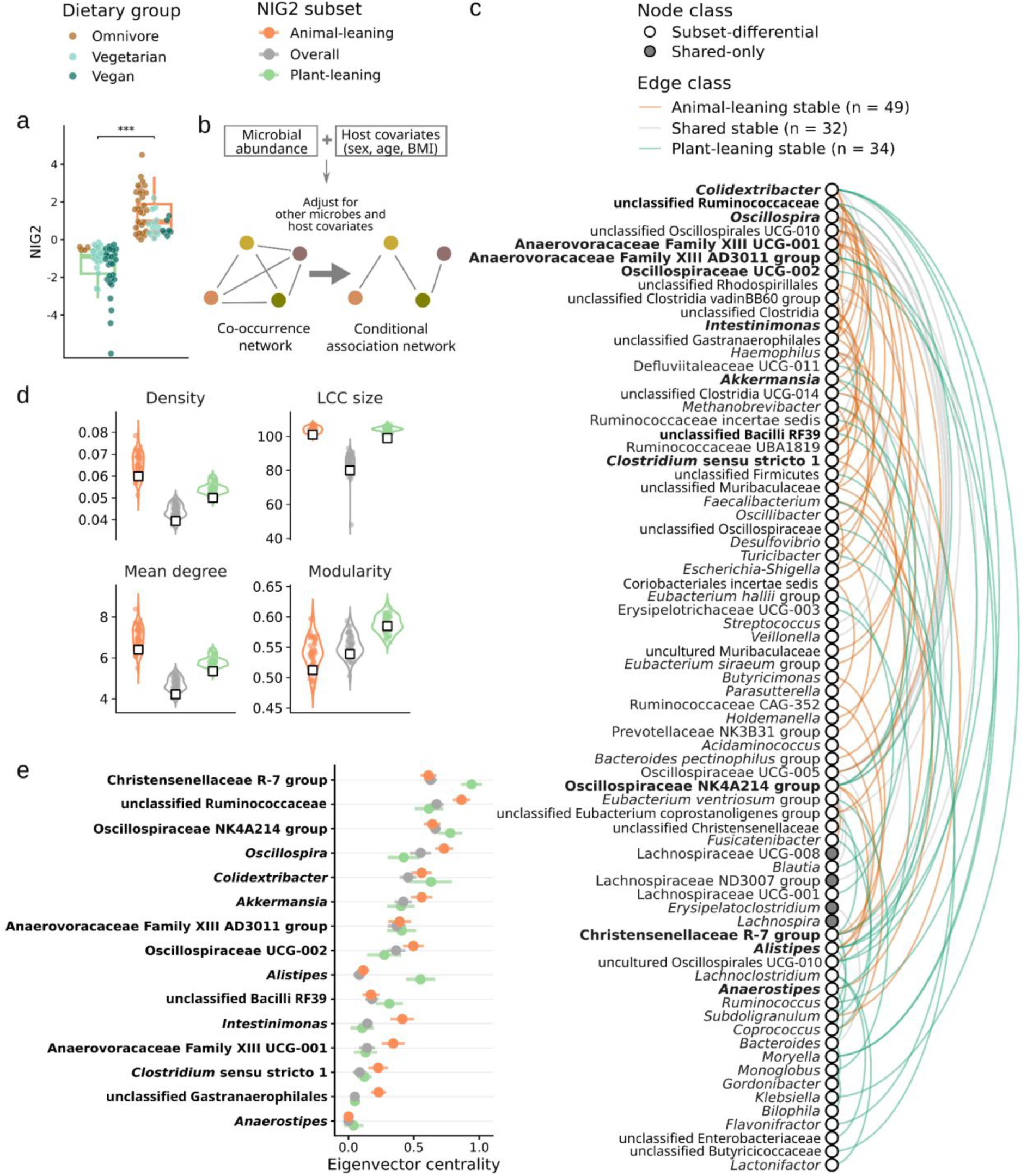
Stability-defined microbial association patterns differ between NIG2-defined animal- and plant-leaning subsets. **a**, Distribution of NIG2 scores used to define animal-leaning and plant-leaning subsets. Participants were stratified by the median NIG2 value, with values above the median classified as animal-leaning and values below the median classified as plant-leaning. Points are colored by dietary groups. **b**, Conditional associations were estimated using a Gaussian graphical model. In contrast to co-occurrence networks based on marginal correlations, conditional association networks retain relationships that remain after accounting for other microbial features and host covariates (sex, age, and BMI). **c**, Linear network plot showing high-confidence microbial associations with differential stability between animal-leaning and plant-leaning subset networks. Edges were classified according to their frequency across repeated subsampling networks. Animal-leaning stable edges were present in ≥90% of animal-leaning subsampling networks and ≤10% of plant-leaning subsampling networks, whereas plant-leaning stable edges were defined conversely. Shared stable edges were present in ≥90% of subsampling networks in both groups. Node class indicates whether a taxon participated in at least one subset-differential stable edge or only in shared stable edges. Bold labels indicate the 15 taxa most frequently involved in subset-differential stable edges. **d**, Topological properties of microbe-only subnetworks inferred after metadata-adjusted network inference. Violin outlines and points show distributions across repeated subsampling networks, and white squares indicate estimates from the full-data networks. LCC denotes the largest connected component. **e**, Eigenvector centrality of the 15 taxa most frequently involved in subset-differential stable edges. Points indicate mean eigenvector centrality across repeated subsampling networks, and horizontal bars indicate ±1 SD.

Stability-based edge comparison revealed high-confidence microbial associations in the two NIG2-defined subsets (Fig. 5c). We identified 49 animal-leaning stable microbial associations, 34 plant-leaning stable microbial associations, and 32 shared stable microbial associations that were consistently recovered across repeated subsampling networks. Taxa most frequently involved in subset-differential stable microbial associations included members of Oscillospiraceae, Ruminococcaceae, Christensenellaceae, and related anaerobic gut-associated genera, suggesting that animal- and plant-leaning nutrient profiles were associated with different stable microbial association patterns.

Topological comparison of microbe-only subnetworks showed that animal-leaning networks tended to have higher density and mean degree than plant-leaning networks, whereas plant-leaning networks showed consistently large LCC size and higher modularity estimates across subsampling networks (Fig. 5d). These patterns suggest that animal- and plant-leaning nutrient profiles were associated not only with differences in individual stable microbial associations, but also with differences in overall community organization.

Among the 15 taxa most frequently involved in subset-differential stable microbial associations, eigenvector centrality showed distinct patterns across the animal-leaning, plant-leaning, and overall networks (Fig. 5e). Several taxa, including Christensenellaceae R-7 group, unclassified Ruminococcaceae, Oscillospiraceae NK4A214 group, and *Colidextribacter*, showed higher centrality in one subset network than in the other. These patterns indicate that taxa frequently involved in subset-differential stable edges can also differ in their relative network prominence, despite showing broadly comparable prevalence and relative abundance between animal- and plant-leaning subsets (Supplementary Fig. S8). Overall, these findings suggest that the continuous animal- to plant-leaning nutrient intake gradient was associated with distinct microbial conditional association structures.

## Discussion

This study shows that a continuous nutrient-derived dietary gradient provides a quantitative metric for characterizing dietary exposure and investigating diet-microbiome-host physiology relationships in healthy adults. By capturing a spectrum from animal- to plant-leaning nutrient profiles, this gradient revealed associations with gut microbiome composition, candidate microbiome-mediated diet-biomarker relationships, and differences in microbial association network structure. These findings indicate that how dietary exposure is quantified can influence the detection and interpretation of diet-microbiome associations.

From an exposome perspective, which aims to characterize the totality of non-genetic environmental exposures influencing health and disease^40,41^, diet represents a major and modifiable exposure, while its accurate quantification remains challenging. Dietary exposure is commonly assessed using food frequency questionnaires, 24-hour dietary recalls, or weighed food records, which estimate dietary intake with different temporal resolution, participant burden, and measurement precision^42^. These data can then be translated into different exposure metrics, ranging from dietary groups such as omnivore, vegetarian, and vegan diets, to predefined dietary indices such as DASH adherence scores^43^, and nutrient-derived dietary patterns. In the present study, dietary exposure was quantified using four-day weighed food records, allowing detailed estimation of individual nutrient intake before deriving the nutrient intake gradient. NIG2 complemented dietary group labels by summarizing the correlated nutrient structure underlying animal- and plant-leaning diets. This individual-level exposure axis is informative for microbiome studies because gut microbial communities respond to combinations of available dietary substrates^44^. Dietary supplement use is an additional source of nutrient exposure that can decouple micronutrient intake from food-based dietary patterns and may influence gut microbiome features^45,46^. In this cohort, the high concordance between food-derived and total-intake NIG2 suggests that supplement-derived nutrients did not substantially alter the dietary gradient, supporting its robustness as a marker of animal- to plant-leaning nutrient profiles.

NIG2 resolved gut microbiome compositional variation that was not detected when diet was represented by omnivore, vegetarian, and vegan dietary groups. This supports the value of nutrient-derived gradients for identifying diet-associated microbiome variation, particularly when sample size limits the resolution of categorical group comparisons. No association was observed between NIG2 and within-sample diversity. This emphasis on community-level differences is broadly consistent with large multi-cohort analyses of omnivore, vegetarian, and vegan populations, in which diet-pattern-specific microbiome profiles were detectable, whereas alpha-diversity differences were not consistently observed^3^. The variation explained by NIG2 was in line with effect sizes typically reported for dietary exposures in population-based microbiome studies^47,48^. The taxa aligned with the NIG2-associated compositional axis provided descriptive taxonomic context for interpreting the animal- to plant-leaning nutrient intake gradient. In particular, the opposite positioning of *Prevotella* 9 and *Bacteroides* was consistent with previously described diet-associated microbiome configurations, with *Prevotella*-rich communities often linked to carbohydrate- and fiber-associated diets^49,50^ whereas higher *Bacteroides* abundance has been reported in animal-based dietary interventions^49,51^. *Blautia* was also located toward the animal-leaning side of the ordination, providing compositional context for its recurrence in the candidate mediation results discussed below. In parallel, taxa oriented mainly along PCoA2, including the Christensenellaceae R-7 group and unclassified Ruminococcaceae, suggested additional structure in microbiome composition beyond the primary NIG2-associated axis.

The mediation analysis complemented the compositional microbiome findings by prioritizing candidate microbiome-mediated diet-biomarker relationships along the animal- to plant-leaning nutrient intake gradient. The estimated mediated effects point to specific microbial genera that may contribute to the observed relationship between NIG2 and host biomarker variation. These candidate relationships spanned thyroid-related, hematological, and vitamin-related biomarkers, all of which are connected to dietary intake or nutritional status. Thyroid-related markers reflect broader nutritional and metabolic status^52^, MCV is a hematological index relevant to vitamin B12 and folate status^53^, and vitamin E and vitamin K1 are fat-soluble vitamins influenced by dietary intake^54,55^. A notable example was the NIG2-vitamin E relationship, for which *Bacteroides* and *Blautia* showed positive mediated effects. This finding is particularly interesting because dietary vitamin E intake was negatively associated with NIG2, whereas circulating vitamin E was positively associated with NIG2 in pairwise analyses. Meanwhile, both genera were positively associated with NIG2. These findings suggest that the NIG2-associated differences in circulating vitamin E may not simply reflect vitamin E intake, but also involve microbial features associated with the animal- to plant-leaning nutrient intake gradient. *Bacteroides* provides a biologically plausible candidate mediator, given its reported associations with dietary fat intake and bile acid metabolism ^56^. Since circulating vitamin E reflects complex processes of absorption, transport, and metabolism rather than dietary intake alone ^57^, the positive mediated effect through *Bacteroides* may indicate that this genus accounts for part of the observed NIG2-vitamin E relationship, although it does not establish a specific mechanistic route. *Blautia* also showed a positive mediated effect for the NIG2-vitamin E relationship, but a direct connection between *Blautia* and circulating vitamin E is not well established. This finding therefore supports its interpretation as a candidate mediator requiring further validation. These mediation results should be interpreted as hypothesis-generating because the cross-sectional design does not establish causality or temporal directionality. Nevertheless, integrating nutrient-derived dietary gradients with microbial and host biomarker data helped prioritize biologically interpretable hypotheses about microbial taxa that may connect dietary exposure with host physiological variation.

Most previous studies have characterized dietary associations through differences in relative abundance and community composition^51,58^. Building on the view of the gut microbiome as an interconnected ecological community^59^, network-based analysis provides an additional perspective by asking whether nutrient-associated dietary variation is also reflected in the conditional association structure among microbial taxa. By estimating conditional rather than simple pairwise correlations^60^, the metadata-adjusted Gaussian graphical model used here reduces the influence of indirect correlations and provides a more conservative view of microbial association structure. The subset-specific stable taxon-taxon associations observed here suggest that NIG2-defined animal- and plant-leaning subsets may differ in their conditional association structure. The topological patterns further suggest that microbial association architecture differed between the two subsets, with the animal-leaning subnetwork showing a more densely connected structure and the plant-leaning subnetwork showing a more modular organization. These results point to two levels of diet-associated network variation: which microbial taxa are linked by stable associations, and how these associations are organized at the community level.

The network analysis also highlighted taxa whose relevance was not apparent from the primary NIG2-associated compositional axis. Christensenellaceae R-7 group and unclassified Ruminococcaceae were not the main taxa aligned with NIG2-associated compositional axis, but they emerged as network-relevant taxa within subset-specific microbial association structures. Christensenellaceae R-7 group showed greater network prominence in the plant-leaning subnetwork, whereas unclassified Ruminococcaceae was more prominent in the animal-leaning subnetwork. This pattern suggests that taxa contributing to additional compositional variation may also occupy different positions within animal- and plant-leaning microbial association structures. *Christensenellaceae* has been linked to lower serum lipid level, healthy BMI, and co-occurrence with other heritable gut microbes^61–63^. Ruminococcaceae includes anaerobic gut taxa involved in complex carbohydrate degradation and fermentation, with members contributing to resistant starch degradation and downstream short-chain fatty acid production^64,65^. The involvement of Christensenellaceae R-7 group and unclassified Ruminococcaceae in subset-specific network patterns suggests that nutrient-related dietary variation may be reflected in the network positioning of metabolism-related gut taxa. These inferred networks therefore provide a structured view of microbial conditional associations beyond relative abundance or community composition alone, while not directly resolving ecological interactions or temporal dynamics.

## Conclusions

In conclusion, this study demonstrates that nutrient-derived dietary gradients provide a useful framework for investigating diet-microbiome relationships beyond categorical dietary groups. The animal- to plant-leaning nutrient intake gradient identified here was associated with gut microbiome composition, candidate microbiome-mediated diet-biomarker relationships, and differences in microbial conditional association structure, suggesting that dietary variation may be reflected across multiple levels of microbiome organization. By integrating nutrient intake, microbiome profiles, host biomarkers, and network analysis within a well-characterized healthy cohort, this work highlights complementary perspectives on how dietary intake patterns are associated with the gut ecosystem. Future studies with larger sample sizes, longitudinal or dietary intervention designs, and complementary multi-omics approaches such as shotgun metagenomics and metabolomics will be needed to validate these findings and clarify the mechanisms underlying the observed associations.

## Statements

### Ethics statement

This study was approved by the Ethics Committee of the University Medical Center Freiburg, Germany (EK-Freiburg 21-1442) and registered at the German Clinical Trials Register (DRKS00027425).

### Data availability

The 16S rRNA gene sequencing data generated in this study have been deposited in the NCBI Sequence Read Archive under BioProject accession PRJNA1499606 and will be available upon publication. Analysis code, figure-generation scripts, and processed analysis tables required to reproduce the main analyses are archived at: https://github.com/jiyanxu/VVO and will be available upon publication. De-identified metadata used for the analyses are provided with the code repository where participant privacy and consent restrictions permit. Additional data are available from the corresponding author upon reasonable request.

### Author contributions

J.X.: formal analysis, investigation, methodology, visualization, writing – original draft, writing – review and editing. M.R.: investigation, methodology. T.U.P.B.: investigation, methodology. A.M.: investigation, laboratory work, participant scheduling, and study administration. L.H.: conceptualization, study planning, and supervision of vitamin B12-related biomarker measurements. R.H.: conceptualization, study planning, study supervision, investigation, and funding acquisition. M.J.: resources, project administration, and writing – review. V.H.J.-D.: supervision of data analysis, methodology, interpretation, and writing – review and editing. S.K.F.-S.: supervision of data analysis, methodology, interpretation, and writing – review and editing. All authors reviewed and approved the final version of the manuscript. M.A.S.: conceptualization, study planning, study supervision, investigation, and funding acquisition.

## Acknowledgments

We thank all study participants for their participation. We also thank the clinical and laboratory staff involved in participant coordination, sample collection, sample processing, and data generation. We acknowledge the support of the Max Cluster high-performance computing infrastructure for computational analyses.

## Generative AI statement

During the preparation of the first draft of this manuscript, the author J.X. used ChatGPT (GPT-5.5) to support English language editing and improve clarity. The authors J.X., V.H.J.-D., and S.K.F.-S. reviewed and edited the AI-assisted text. All authors read and approved the final version of the manuscript and take full responsibility for the content of it.

