## Supplementary Figure for "A nutrient-derived dietary gradient links gut microbiome composition, network structure, and host physiology"

### Supplementary materials

Supplementary Fig. S1 | Hierarchical clustering of blood biomarkers and selection of representative biomarkers for downstream analyses
Supplementary Fig. S2 | Distribution of the nutrient intake gradient (NIG2)
Supplementary Fig. S3 | Principal component analysis of nutrient intake and characterization of nutrient intake gradients
Supplementary Fig. S4 | Associations between nutrient intake gradients and demographic covariates
Supplementary Fig. S5 | Sensitivity analysis of nutrient intake gradients including supplement-derived intake
Supplementary Fig. S6 | Associations between NIG2 and gut microbiome alpha diversity
Supplementary Fig. S7 | Associations between dietary group and gut microbiome composition

Supplementary Table S1. Blood biomarker information and representative biomarker selection for downstream analyses
Supplementary Table S2. Nutrient variables, scaling parameters, and PCA loadings used to define food-derived and total-intake nutrient intake gradients
Supplementary Table S3. Forward and reverse-direction mediation results

Supplementary Table S4. Abbreviations and labels used in the study

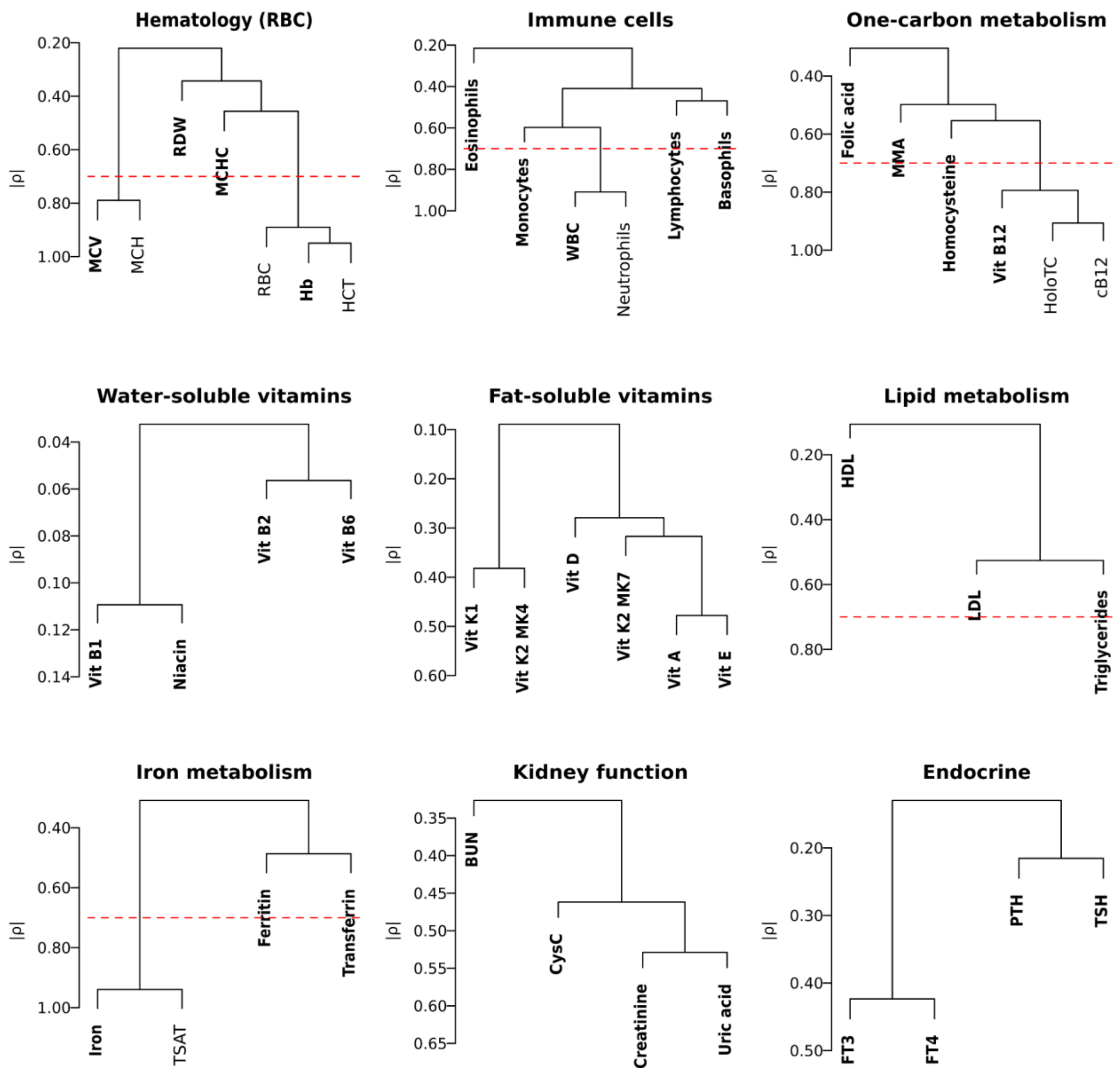


**Supplementary Fig. S1: Hierarchical clustering of blood biomarkers and selection of representative biomarkers for downstream analyses.** Blood biomarkers were hierarchically clustered within biologically defined categories using average-linkage hierarchical clustering based on the distance metric $1 - |\rho|$, where $\rho$ denotes the pairwise Spearman correlation coefficient. The red dashed line indicates the clustering threshold corresponding to $|\rho| = 0.7$. Biomarkers retained as representative variables for downstream analyses are shown in bold. Liver function biomarkers are not shown because only two liver function biomarkers (GPT and GOT) were available in this cohort. Their pairwise correlation was $|\rho| = 0.728$, and GPT was selected as the representative biomarker because it is a more liver-specific marker of hepatocellular injury.

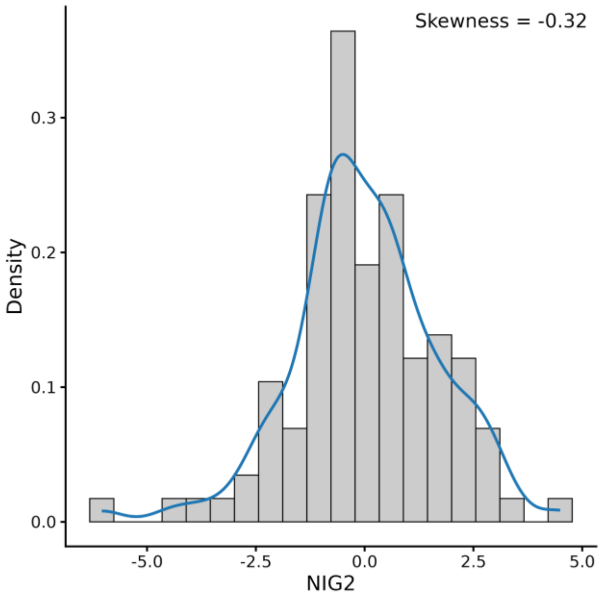


**Supplementary Fig. S2: Distribution of the nutrient intake gradient (NIG2).** Histogram and kernel density estimate of the nutrient intake gradient (NIG2) across all participants. The distribution showed no pronounced skewness (skewness = −0.32), supporting the use of the original NIG2 values without log-transformation prior to standardization and downstream analyses.

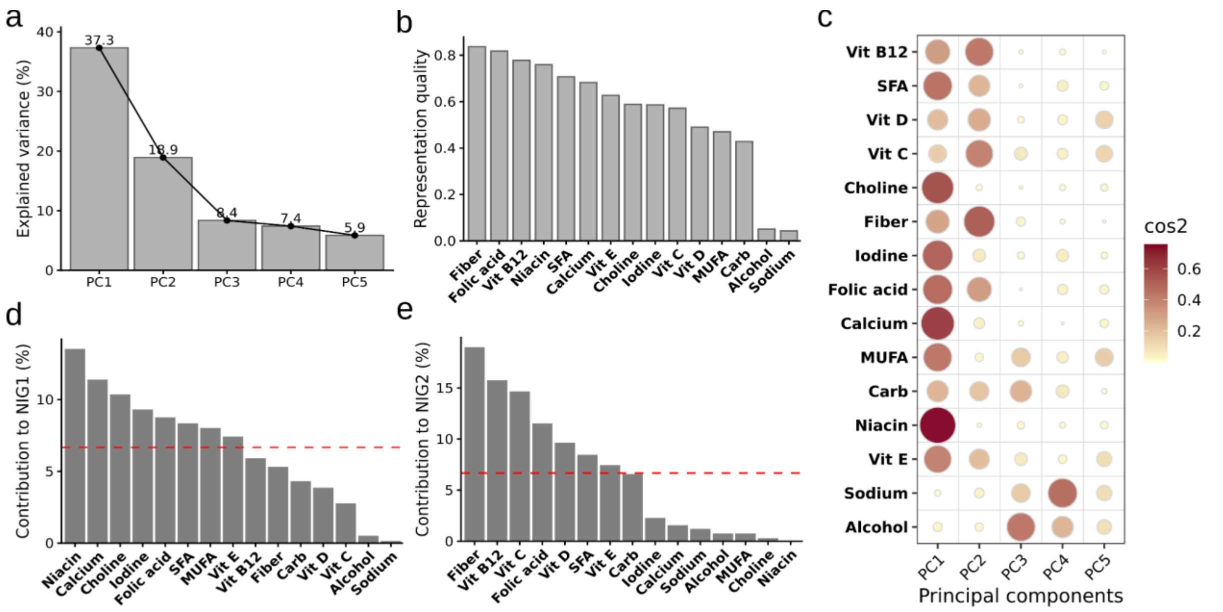

**Supplementary Fig. S3: Principal component analysis of nutrient intake and characterization of nutrient intake gradients. a,** Scree plot showing the percentage of variance explained by the first five principal components. The first two principal components were retained and are hereafter referred to as nutrient intake gradient 1 (NIG1) and nutrient intake gradient 2 (NIG2). **b,** Nutrient representation quality on the NIG1-NIG2 plane, quantified as the cumulative squared cosine ($\cos^2$) across both axes. Higher values indicate that a nutrient is better represented by the two-dimensional PCA plane. **c,** Bubble plot showing nutrient-specific $\cos^2$ values across the first five principal components. **d,e,** Nutrient contributions to NIG1 (**d**) and NIG2 (**e**). Dashed red lines indicate the expected average contribution under equal contribution of all nutrients.

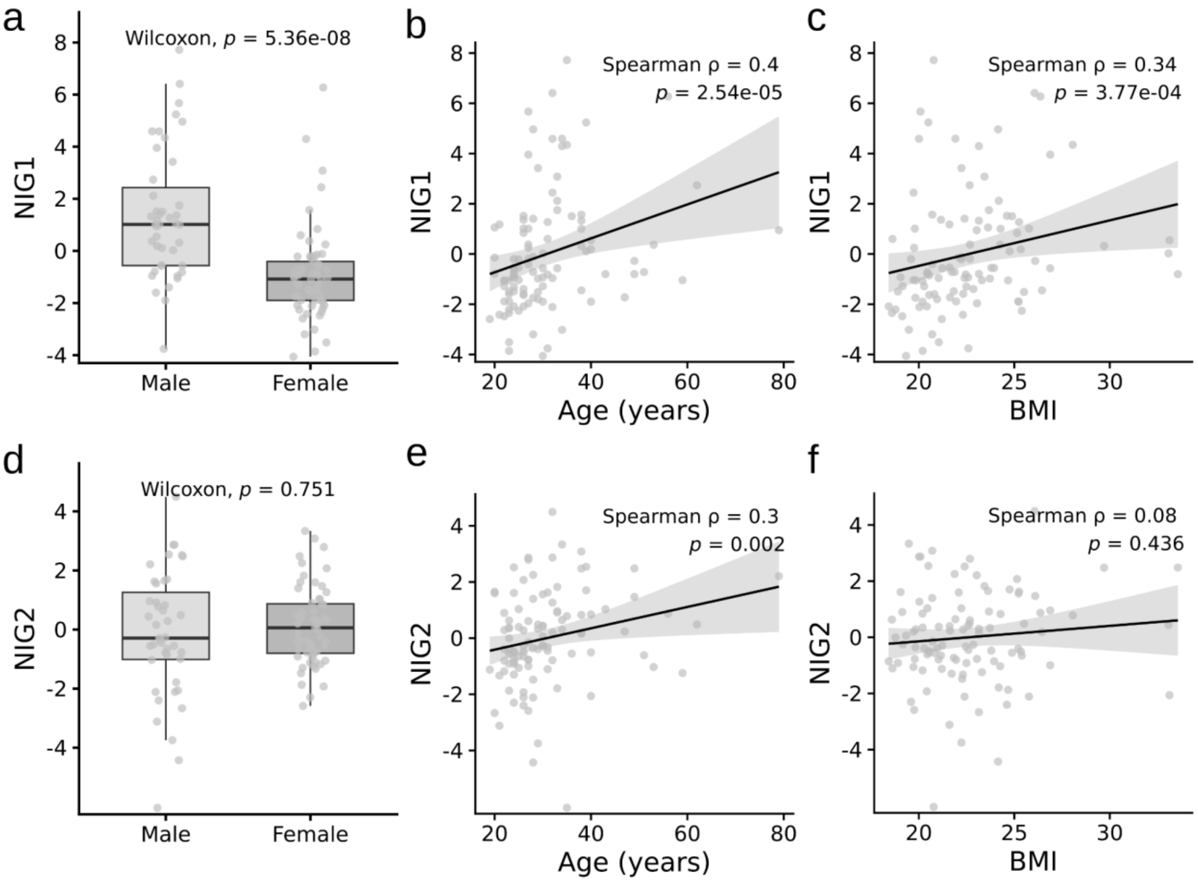

**Supplementary Fig. S4: Associations between nutrient intake gradients and demographic covariates.** Associations of NIG1 (**a–c**) and NIG2 (**d–f**) with sex, age, and BMI. Sex differences were assessed using Wilcoxon rank-sum tests, whereas associations with age and BMI were assessed using Spearman correlation. Scatter plots show individual participants, fitted linear trends, and 95% confidence intervals. NIG1 was associated with sex, age, and BMI, whereas NIG2 was associated with age but not with sex or BMI.

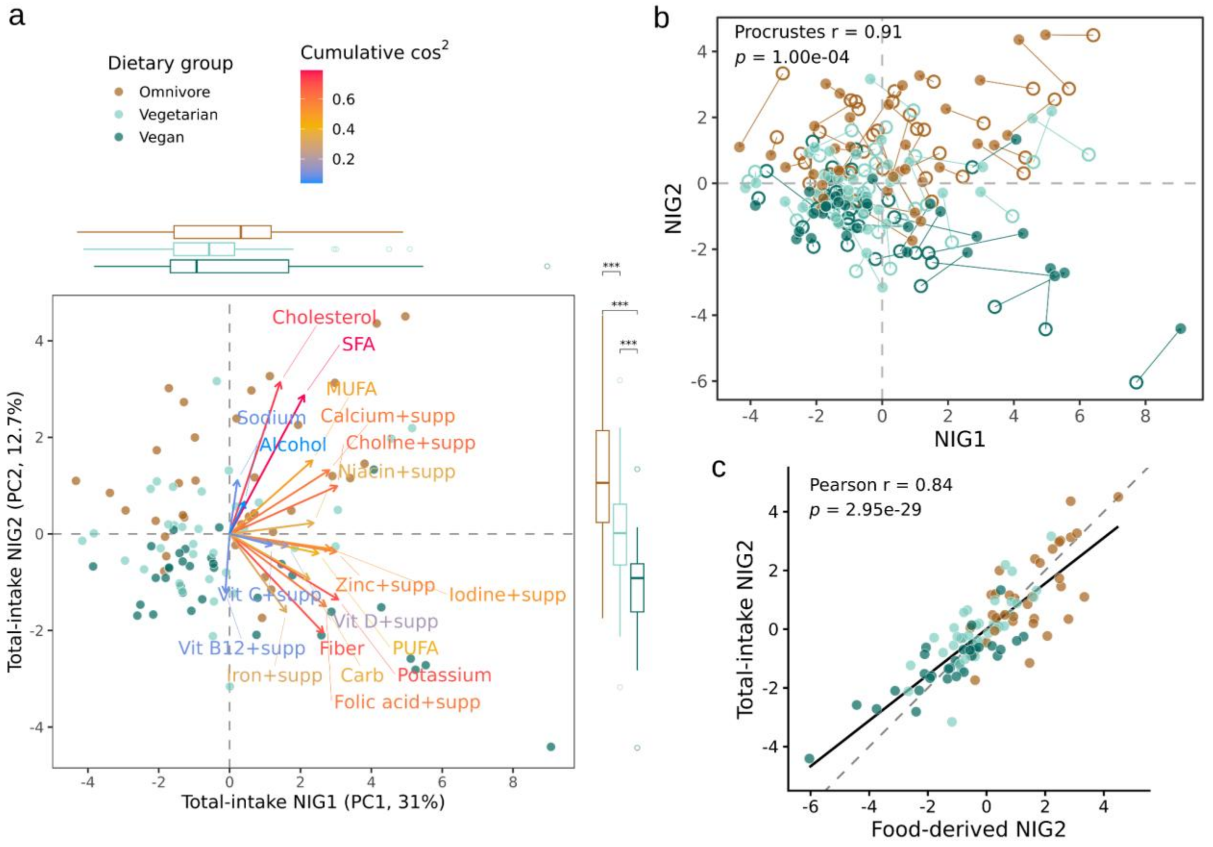

**Supplementary Fig. S5: Sensitivity analysis of nutrient intake gradients including supplement-derived intake. a**, Principal component analysis (PCA) was repeated using total nutrient intake, including food-derived intake and supplement-derived intake where available. Each point represents an individual participant and is colored by dietary group. Nutrient arrows indicate loading vectors on the first two total-intake principal components, referred to as total-intake NIG1 and total-intake NIG2. Arrow color denotes the cumulative cos² across the first two components. Marginal boxplots show the distribution of total-intake NIG1 and total-intake NIG2 scores across dietary groups. Statistical significance in marginal boxplots was assessed using one-way ANOVA followed by Tukey's honestly significant difference post hoc test. ***Adjusted $p < 0.001$. Nutrients labeled "+supp" indicate total intake variables that include supplement-derived intake in addition to food-derived intake. **b**, Procrustes comparison of food-derived and total-intake PCA configurations on the first two nutrient intake gradients. Open circles indicate food-derived PCA scores, filled circles indicate total-intake PCA scores, and arrows connect the corresponding positions for each participant after Procrustes alignment. **c**, Pearson correlation between food-derived NIG2 and total-intake NIG2 scores. The dashed line indicates the identity line, and the solid line indicates the fitted linear regression line.

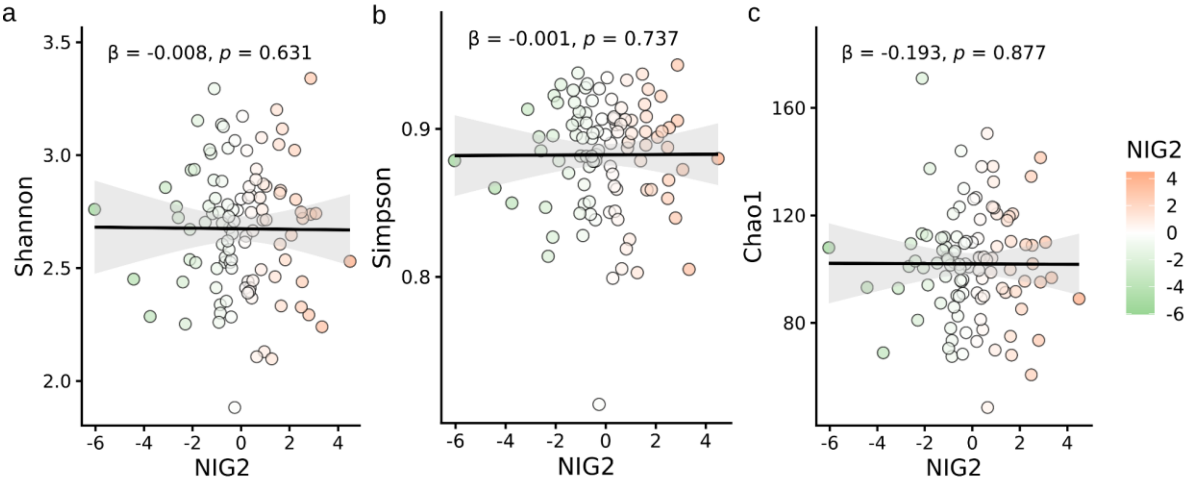


**Supplementary Fig. S6: Associations between NIG2 and gut microbiome alpha diversity.** Associations of NIG2 with Shannon, Simpson, and Chao1 alpha diversity indices at the genus level. Associations were assessed using linear regression models adjusted for sex, age, and BMI. Points represent individual samples and are colored by NIG2. Solid lines indicate fitted linear regressions, and shaded areas indicate 95% confidence intervals. No significant association was observed between NIG2 and any alpha diversity index.

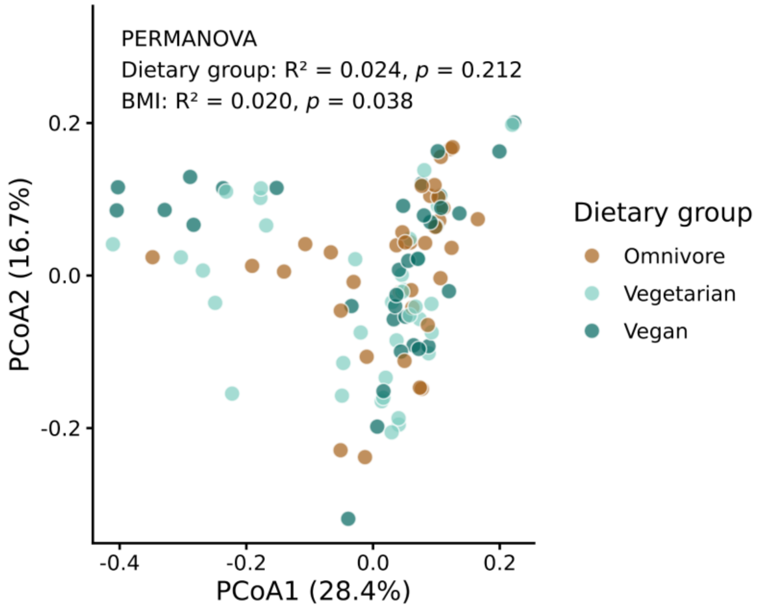


**Supplementary Fig. S7: Associations between dietary group and gut microbiome composition.** Principal coordinates analysis (PCoA) of Bray-Curtis dissimilarities calculated from genus-level microbial profiles. Samples are colored according to dietary group (omnivore, vegetarian, and vegan). PERMANOVA adjusted for sex, age, and BMI showed no significant association between dietary group and overall microbial community composition ($R^2 = 0.024$, $p = 0.212$).

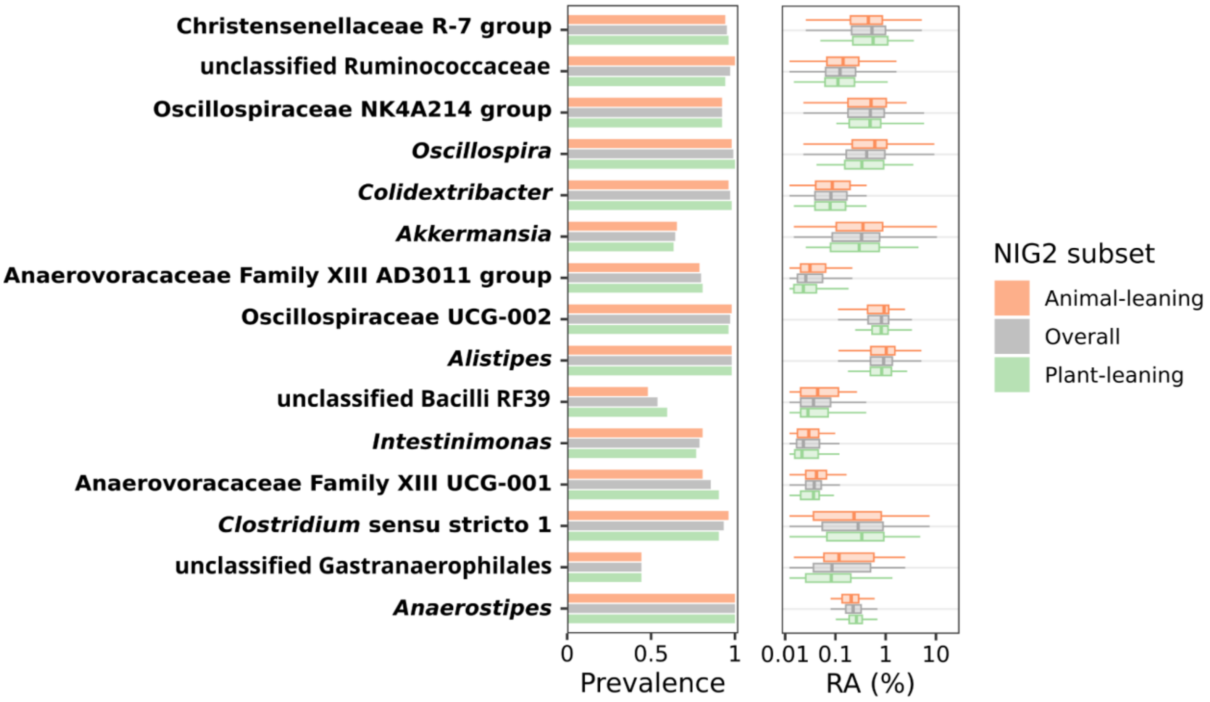

**Supplementary Fig. S8: Prevalence and relative abundance of taxa most frequently involved in subset-differential stable microbial associations.** Prevalence and relative abundance of the 15 taxa most frequently involved in subset-differential stable edges between NIG2-defined animal- and plant-leaning microbial association networks. Taxa are shown in the same order as in the eigenvector centrality plot in Fig. 5e. The left panel shows prevalence. The right panel shows genus-level relative abundance (RA, %) displayed on a log10-scaled axis. Overall represents all samples and is shown for descriptive reference only. Fisher's exact tests showed no significant differences between animal- and plant-leaning subsets in prevalence after Benjamini–Hochberg correction. Linear models adjusted for sex, age, and BMI showed no significant differences in relative abundance between animal- and plant-leaning subsets for these taxa after Benjamini–Hochberg correction.
